# Phylogeographic model selection using convolutional neural networks

**DOI:** 10.1101/2020.09.11.291856

**Authors:** Emanuel Masiero da Fonseca, Guarino R. Colli, Fernanda P. Werneck, Bryan C. Carstens

## Abstract

The field of phylogeography has evolved rapidly in terms of the analytical toolkit to analyze the ever-increasing amounts of genomic data. Despite substantial advances, researchers have not fully explored all potential analytical tools to tackle the challenge posed by the huge size of genomic datasets. For example, deep learning techniques, such as convolutional neural networks (CNNs), widely employed in image and video classification, are largely unexplored for phylogeographic model selection. In non-model organisms, the lack of information about their ecology, natural history, and evolution can lead to uncertainty about which set of demographic models should be considered. Here we investigate the utility of CNNs for assessing a large number of competing phylogeographic models using South American lizards as an example, and approximate Bayesian computation (ABC) to contrast the performance of CNNs. First, we evaluated three demographic scenarios (constant, expansion, and bottleneck) for each of four recovered lineages and found that the overall model accuracy was higher than 98% for all lineages. Next, we evaluated a set of 26 models that accounted for evolutionary relationships, gene flow, and changes in effective population size among these lineages and recovered an overall accuracy of 87%. In contrast, ABC was unable to single out a best fit model among 26 competing models. Finally, we used the CNN model to investigate the evolutionary history of two South American lizards. Our results indicate the presence of hidden genetic diversity, gene flow between non-sister populations, and changes in effective population sizes through time, likely in response to Pleistocene climatic oscillations. Our results demonstrate that CNNs can be easily and usefully incorporated into the phylogeographer’s toolkit.

## Introduction

One key research goal of phylogeographic research has been to investigate how historical processes have shaped genetic variation across geographic space. In the early years of phylogeography, interpretations were highly qualitative and largely based on gene genealogies and the geographic distribution of the haplotypes. Because of their descriptive nature, such phylogeographic investigations were susceptible to overinterpretation (Knowles & Maddison, 2002), where a detailed explanation of the causes of intraspecific diversification usually went beyond the evidence supported by the data, and confirmation bias (Nickerson, 1998), where researchers often interpreted new results in a manner that supported previous findings (Carstens et al., 2009). As the field matured, researchers recognized the importance of statistical approaches that explicitly incorporate uncertainty to draw meaningful conclusions about species’ evolutionary history. Therefore, the identification of statistical models relevant for data analysis is a crucial step of any model-based phylogeographical investigation.

Phylogeographers have employed three general approaches to identify the models used to describe the data and make inference: (i) intuitive model identification; (ii) phylogeographic hypothesis testing; and (iii) objective model selection (Carstens et al., 2017). In the first approach, researchers use a particular evolutionary model to estimate a set of parameters of interest based on their expertise about the organism and its environment. Although this approach has enabled the evaluation of complex evolutionary processes, it can lead to unreliable estimates of the parameters of interest due to model misspecification (Koopman & Carstens, 2010). Biological intuition often drives the choice of the analytical framework(s) used to analyze the data. For example, researchers may choose to analyze their data with an isolation with migration model or an *n*-island migration model due to beliefs regarding the processes that have influenced their system. In practice, if the chosen model has a poor fit to the evolutionary history of the organism, the resulting inferences can be misleading (Beerli & Palczewski, 2010; Hey et al., 2015). Unfortunately, the estimation of many evolutionary processes eventually becomes intractable in a likelihood framework (Beaumont, 2010; Beaumont et al., 2002), Therefore, no single analytical method can incorporate all possible evolutionary processes and use maximum likelihood or Bayesian methods to identify parameter values that maximize the probability of the model given the data. Hypothesis testing (e.g., Knowles et al., 2007) is conducted under an assumed model and, thus, subject to the same potential flaws as intuitive approaches. For these reasons, many researchers now utilize model selection approaches in phylogeographic research.

Simulation-based and likelihood-free approaches, which can accommodate complex demographic scenarios (Pritchard et al., 1999), are often used by researchers to conduct phylogeographic model selection. Software such as *ms* (Hudson, 2002), *msprime* (Kelleher et al., 2016), and *fastsimcoal2* (Excoffier et al., 2013) can be used to conduct coalescent simulations under customized demographic models that can approximate the details of almost any empirical system. After the simulation procedure, empirical and simulated datasets can be statistically evaluated using a variety of methods, including hypothesis testing (e.g., Knowles et al., 2007), Approximate Bayesian Computation (ABC; e.g., Fagundes et al., 2007), information theory (e.g., Carstens et al., 2009), and machine learning approaches such as Random Forest (Smith et al., 2017). While these have in common the flexibility to assess multiple demographic models given the observed data, factors such as the type of data collected and details about the empirical system make it likely that there isn’t a single “best” approach for all questions.

Information theoretic approaches can be conducted either on SNP data, summarized as site frequency spectra (SFS; e.g., Thomé & Carstens, 2016), or gene trees (e.g., Jackson et al., 2017). Such approaches are effective at considering large numbers of models, but at the expense of parameter estimation. Approximate Bayesian Computation (ABC) remains a widely used approach in demographic model selection, but can potentially suffer from the “curse of dimensionality” when comparing more than a handful of demographic models (Pelletier & Carstens, 2014; Schrider & Kern, 2018). The computational effort of these approaches varies, but ABC becomes computationally expensive when the data are summarized on a locus by locus basis. For this reason, methods that summarize SNP data as SFS and use machine learning to identify the best model are increasingly being applied (e.g., Pudlo et al., 2016; Smith et al., 2017). As genomic data become easier to collect and more common in non-model systems, increased exploration of the usefulness of these (and other) approaches to phylogeographic model selection is warranted.

### Supervised Machine Learning

Supervised machine learning (SML) is a branch of artificial intelligence that gives computers the ability to learn from data without being explicitly programmed and where labels (i.e., pre-classified data) are available for all the samples. SML involves (i) training a predictive model using a subset of a labeled dataset, (ii) evaluating the model using the remaining portion of the labeled dataset, and (iii) using the now-trained model to predict new, unlabeled examples. One example of a SML approach to phylogeographic inferences is implemented in the R package delimitR (Smith & Carstens, 2020), which uses a Random Forest classifier to create hundreds of individual decision trees (a forest) from SNP data, summarized using SFS, to train the model. Next, the set of decision trees are combined via a consensus tree, and this tree is used to classify a new dataset. Results from a simulation study indicate that delimitR is able to compare hundreds of alternative models with high accuracy, even when comparing complex evolutionary scenarios. However, results in other fields that apply SML approaches indicate that Random Forest may not be as efficient as other approaches, such as convolutional neural networks (CNN; Box 1; Razzak et al., 2018). Since CNNs take as input a set of labeled images and train a model to predict the content of new images, one potential advantage of this approach is that data do not need to be summarized using standard genetic summary statistics or a SFS. Rather, prediction can be made directly from the alignment containing the genetic variation from sampled individuals (Flagel et al., 2019). CNNs have been used to address a range of biological questions, from detecting selective sweeps (Flagel et al., 2019) to predicting cancer outcomes (Mobadersany et al., 2018). In spite of all its benefits, the potential applicability of CNNs to phylogeographic model selection remains largely unexplored.

Here we explore the usefulness of CNNs for phylogeographic model selection. We use a simulation-based approach to create labeled examples (i.e., DNA alignments), converted to a black and white image by labeling the major allele as the ancestral state and the minor allele as a derived state. After training the model using 80% of the labeled data and evaluating its performance using the remaining 20% of the data, we compare the performance of CNNs and ABC to inquire about the evolutionary history of two species of lizards, from contrasting environments in South America.

## Box 1 Overview of Convolutional Neural Networks (CNNs)

Artificial neural networks (ANNs) were proposed as an attempt to mimic the network of neurons that constitute the animal brain. In human brains, for example, an external stimulus is passed through a chain of neurons that culminate in a response. Likewise, ANNs are fed with data (i.e., stimulus) which are passed through an artificial network of neurons to make a prediction (i.e., response). CNNs (also known as ConvNets) are a class of artificial neural networks that use a set of labeled images (input data) to build a model to differentiate among the various labels (e.g., a model able to differentiate between images of cats and dogs). First, the input images (Figure 1a) are transformed into arrays (Figure 1b), and then a convolution operation is performed by multiplying each value in the array by a learnable weight within a kernel (Figure 1c). After the convolution operation, the arrays are converted into a feature map (Figure 1d) where each value is passed through a non-linear function (e.g., ReLU, tanh, sigmoid). Next, a pooling method (maximum, average pooling, etc.) is applied to the feature maps within a kernel to reduce the dimensions of the feature maps and maintain conceivably important features from the convolutional kernel (Figure 1e). These steps can be replicated “n” times inside the CNN architecture. For example, in Figure 1, the convolutional and pooling steps were replicated twice. Lastly, the resulting array of all these operations is flattened into a one-dimensional array and fully connected to an ANN. Together, these steps comprise the forward propagation, in which the goal is to pass the data through the CNN (or ANN) and compute a loss function with respect to the weights. Once the loss function is computed, the CNN works backward (back-propagation) to optimize the weights and minimize the total loss function of the model using partial derivatives. In summary, a set of images is forward propagated into a CNN to calculate a loss function, which in turn is back-propagated to optimize the model weight and minimize the loss function. Thus, the training of a CNN consists of an iterative process of forward and backward propagation. Definitions of commonly used terms in this study are presented in Table 1 and a more detailed description of CNNs is available in Lecun et al. (2015) and Flagel et al. (2019).

**Figure 1.**
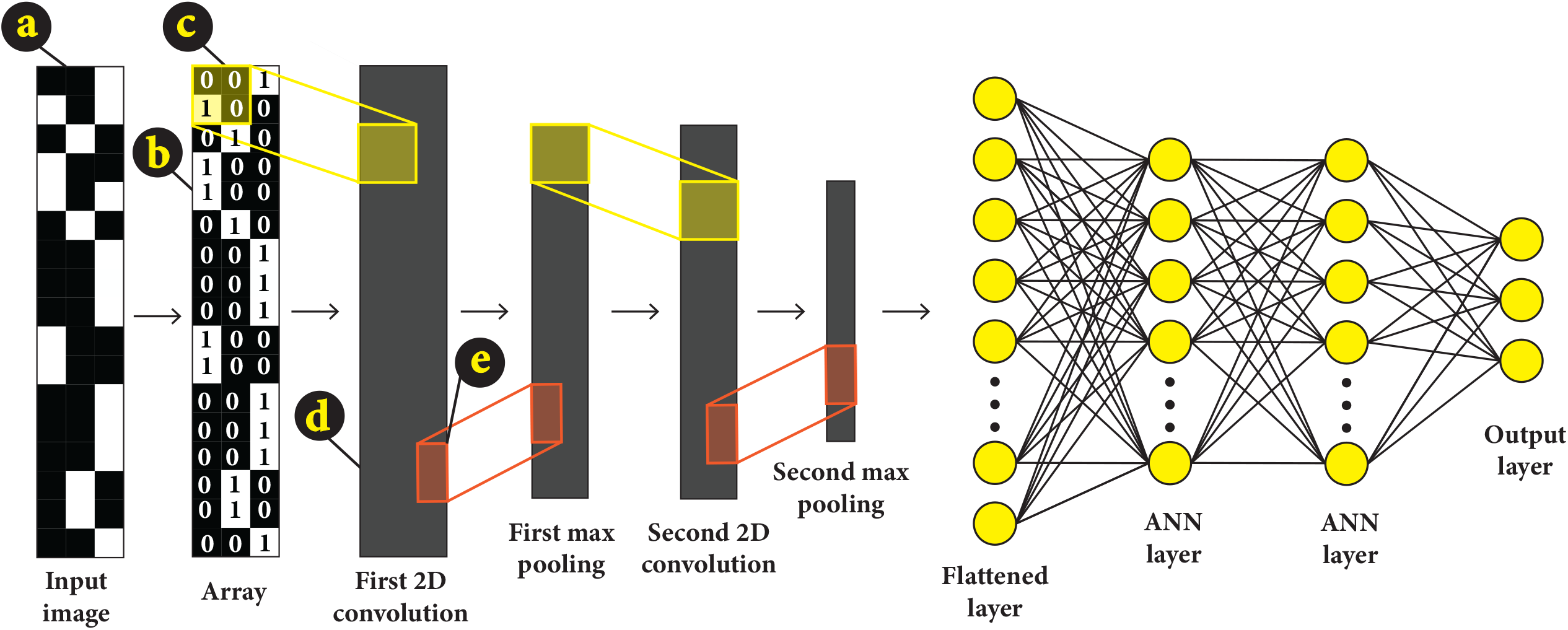
Schematic representation of a 2D convolutional neural network (CNN) architecture. (a) input image; (b) array derived from the input image; (c) convolutional kernel (yellow); (d) feature map; (e) pooling kernel (orange). ANN = artificial neural network.

**Table 1.** A glossary of terms used in this study.

| Term | Definition |
| --- | --- |
| Artificial neural network-ANN | a model of connected layers that attempt to mimic the way that the brain analyzes and processes information |
| Convolutional neural network-CNN | a type of artificial neural network used for image classification and recognition |
| CNN architecture | the general structure of the model that includes the number of convolution and pooling layers, size and numbers kernels, and the number of neurons in each hidden layer. |
| Kernel | vector of weights used for feature detection |
| Neuron | a mathematical function that takes a group of input and weights, applies an activation function (e.g., ReLU, tanh, sigmoid) and output a value |
| Loss function | a method to evaluate how well the model describe the dataset |
| Epoch | the number of times that all images are fed into the model |
| Optimizer | a mathematical function used to update the weights of the model to minimize the loss function |

## Material and Methods

### South American lizards as a case study

We used lizards as a case study to assess the usefulness of CNNs for phylogeographic model selection. Lizards are a diverse group of vertebrates, recognized as model organisms for evolutionary studies due to low thermal tolerance, relatively short generation times, and low dispersal rates (Camargo et al., 2010). For this study, we selected the sister species *Norops brasiliensis* and *N. planiceps* as targets for objective model selection. Little is known about their ecology, natural history, and evolution, which poses great uncertainty about which set of models are appropriate. *Norops brasiliensis* is a terrestrial and diurnal species that occurs predominantly in open areas in the Cerrado and enclaves of Cerrado in Amazonia (Figure 2; Avila-Pires, 1995; Ribeiro, 2015) (Figure 2). While *N. planiceps* is also terrestrial and diurnal, this species is endemic to northern Amazonia, inhabiting mainly “terra firme” forests, which are not periodically flooded (Figure 2; Avila-Pires, 1995; Ribeiro, 2015).

**Figure 2.**
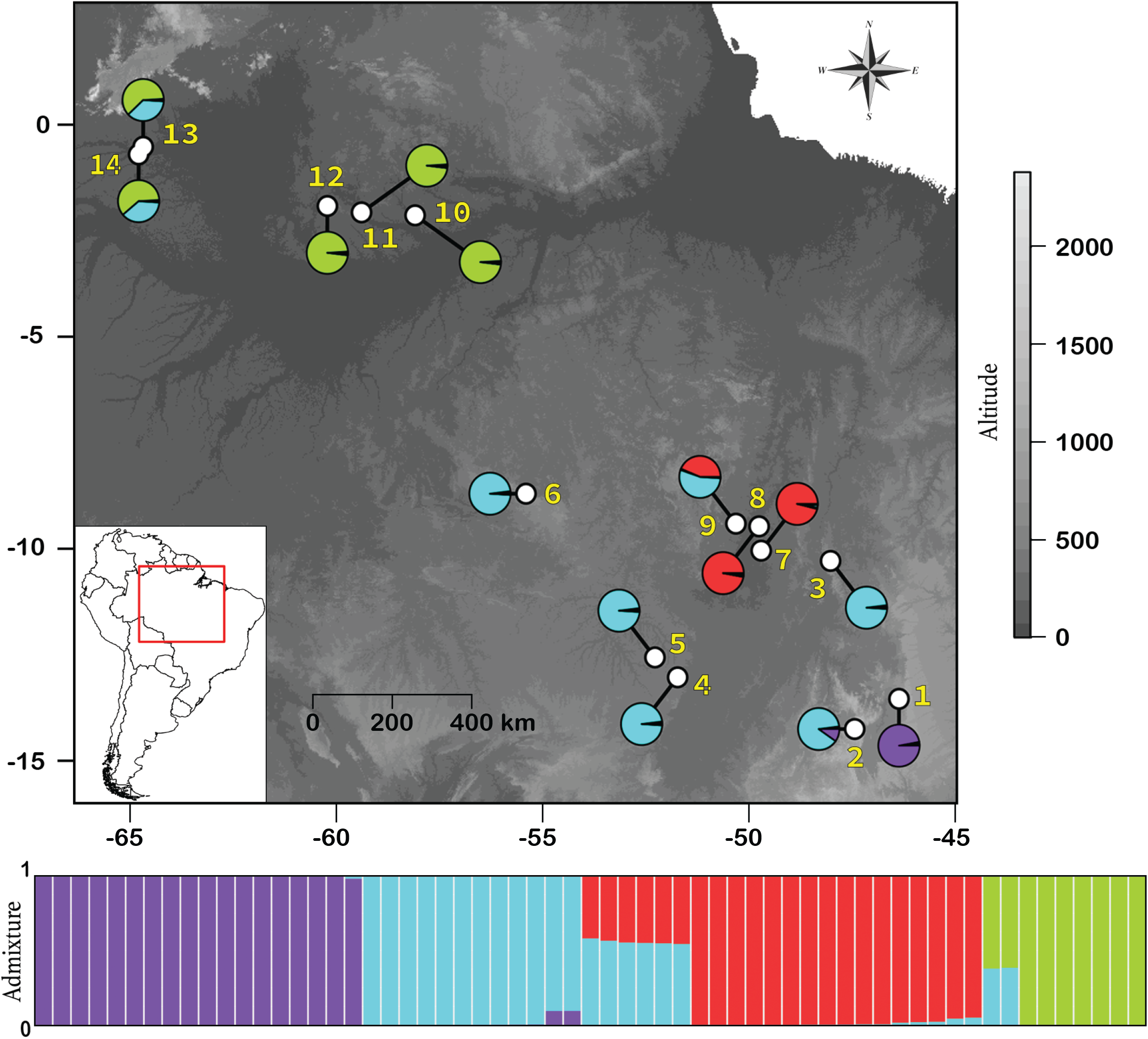
Map showing the geographic distribution of sampled localities. Purple circle = *Norops brasilinesis* (population 1); blue circles = *N. brasilinesis* (population 2); red circles = *N. brasilinesis* (population 3); green circles = *Norops planiceps*. Bar represents the genetic structure of *Norops* ssp. across the area of study according to STRUCTURE analysis.

Amazonia and Cerrado are the largest Brazilian biomes, which together originally covered about 73% of the Brazilian territory. Amazonia is a region predominantly covered by tropical rainforests, whereas the Cerrado is a world hotspot priority for conservation (Myers et al., 2000), characterized by sclerophyllous, fire-adapted flora, abundant grasses and short, thick-barked, and twisted trees (savanna-like vegetation). The Cerrado is part of the South American diagonal of “open formations” (also known as “dry diagonal” or “savanna corridor”) and shares its north-western boundary with Amazonia.

### Sampling and data collection

We obtained 61 tissue samples; 52 from *N. brasiliensis* (nine localities) and 9 from *N. planiceps* (five localities; Figure 1). Samples were obtained from the Herpetological Collection of Brasília University (CHUNB) and the Collections of Amphibians and Reptiles and Genetic Resources from the National Institute of Amazonian Research (INPA-H and INPA-HT).

We extracted DNA from liver or muscle tissues using E.Z.N.A. Tissue DNA Kit and prepared libraries from each species for sequencing using a modified version of the Genotyping-by-Sequencing (GBS) protocol described in Elshire et al. (2011). For DNA digestion, we used 100 ng of freshly extracted DNA and the restriction enzyme Sbf1. After digestion–ligation reactions, we pooled all samples and purified using Agencourt AMPure beads. We amplified samples with polymerase chain reaction (PCR) as follows: (1) initial denaturation at 72 °C for 5 min; (2) 16 cycles consisting of: 98 °C for 10 s for denaturation, 65 °C for 30 s for annealing, and 72 °C for 30 s for extension; (3) final extension at 72 °C for 5 min. Then, we quantified PCR products using the BR DNA Qubit Quantification Kit. To select DNA fragments of 200–500 bp, we used the Blue Pippin Prep and carried out sequencing at the Ohio State University Comprehensive Cancer Center.

### Data processing

We processed (sorted, demultiplexed, clustered, and formatted) raw data from Illumina outputs with ipyrad v 0.9.52 (Eaton & Overcast, 2020), using the resources provided by the Ohio Supercomputer Center. We processed five different datasets: (1) all samples; (2) *N. brasiliensis* (population 1); (3) *N. brasiliensis* (population 2); (4) *N. brasiliensis* (population 3); (5) *N. planiceps*. Datasets 2-5 represent distinct populations recovered in the population assignment analyses (see population assignment section). First, we demultiplexed raw data using individual barcode adapters. Next, we filtered for adapters using the stricter option. We set the maximum low-quality base calls in the read 5, only allowing reads longer than 35 bp. We clustered reads within each sample if their similarity was greater than 85%, set the maximum cluster depth within samples to 10,000 reads, and used a minimum depth for statistical base calling of six reads. Because CNNs do not allow missing data (see CNN section), we removed loci with missing data.

### Population assignments

We used STRUCTURE v2.3.4 (Pritchard et al., 2000) to partition samples into discrete populations before building demographic models. We ran three independent replicates using 100,000 steps of burn-in, followed by 500,000 generations. We performed all runsunder an admixture model for population ancestry and allele frequencies correlated among populations. We evaluated *K*-values ranging from 2 to 6, with ten replications. Using the ad hoc statistic ΔK, we evaluated the optimal value of *K*, calculating the rate of change in the log probability of data between successive *K* values (Evanno et al., 2005), as in STRUCTURE HARVESTER (Earl & vonHoldt, 2012). We combined all replicate analyses under the best value of *K* using the software CLUMPP (Jakobsson & Rosenberg, 2007), and assigned individuals to populations based on their admixture proportion. For example, if an individual was assigned jointly to two populations, we placed that individual in the population with the higher admixture proportion.

### Testing diversification history using convolutional neural networks

In phylogeographic model selection, there are countless ways of parameterizing a model. As the number of lineages and possible parameters increase, the number of possible models grows at a greater than exponential rate. For example, for the four populations we inferred based on the STRUCTURE results, there are more than 2,000 possible models when incorporating topology (four populations), gene flow (isolation vs secondary contact), and changes in population size (constant, bottleneck, and expansion). To facilitate comparison of all potential models, we divided the analysis in two parts. First, we independently tested each population for demographic change in population size through time (12 models). Second, we applied this model of population size change while testing models that consider all possible topologies for four tips and also various migration scenarios (26 models). With this approach, we reduced the model space from more than 2,000 to 38 competing models, which greatly facilitated the comparison between the CNN and ABC approaches to model selection (below).

#### Testing population trajectory through time

In the first part of model selection, we used a CNN to identify the population trajectory that best described the demographic history of each population. We defined three possible scenarios (Figure 2): (a) *constant population size through time*, (b) *population expansion since the last glacial maximum (LGM), and* (c) *population bottleneck since the LGM*. We used the software fastsimcoal2 to simulate 10,000 data examples for each demographic scenario and population. We simulated short DNA sequences (5 bp) for 100,000 independent loci to ensure that the simulator only generated 1 SNP per locus and kept the same number of SNPs as observed in the empirical datasets. We parameterized the ancestral effective population size, current effective population size, and time of population size changing. All priors are presented in Table S1. Next, we wrote custom R scripts to convert the alignment of each simulation into a biallelic matrix, with *n* rows and *k* columns, corresponding to the number of samples and SNPs, respectively. We labeled the major allele as the ancestral state (0) and the minor allele as the derived state (1), such that the matrix could be converted to a black and white image with each entry corresponding to a pixel in the image.

We implemented a two-dimensional CNN architecture as follows: a two-dimensional convolution layer (kernel = 3 ⨯ 1), a two-dimensional maximum pooling layer (kernel = 3 ⨯ 1), a two-dimensional convolutioAn layer (kernel = 3 ⨯ 1), and a two-dimensional maximum pooling layer (kernel = 3 ⨯ 1). We then flattened the output layer from the last pooling. Next, we created a fully connected layer with 100 neurons, followed by one with 25 neurons, and a final layer with three neurons, which correspond to our three demographic models (i.e., constant, expansion, and bottleneck; Figure 3). For all layers, we used rectified linear unit activation functions (ReLU), except for the last one where we used a softmax function. This function is a generalization of the logistic function and used for multiclass prediction. We compiled the CNN using the Adam optimization procedure (Kingma & Ba, 2015), a categorical cross-entropy loss function, and a mini-batch size of 100. We ran the CNN for 10 epochs, although without any improvement after three epochs. We did not include a dropout layer because of the lack of evidence of overfitting. We trained the CNN using 80% of the simulated datasets and used the remaining 20% to evaluate model accuracy. Lastly, we used the trained model to predict the model that likely generated the empirical dataset. We built all CNNs with the Keras python library (https://keras.io).

**Figure 3.**
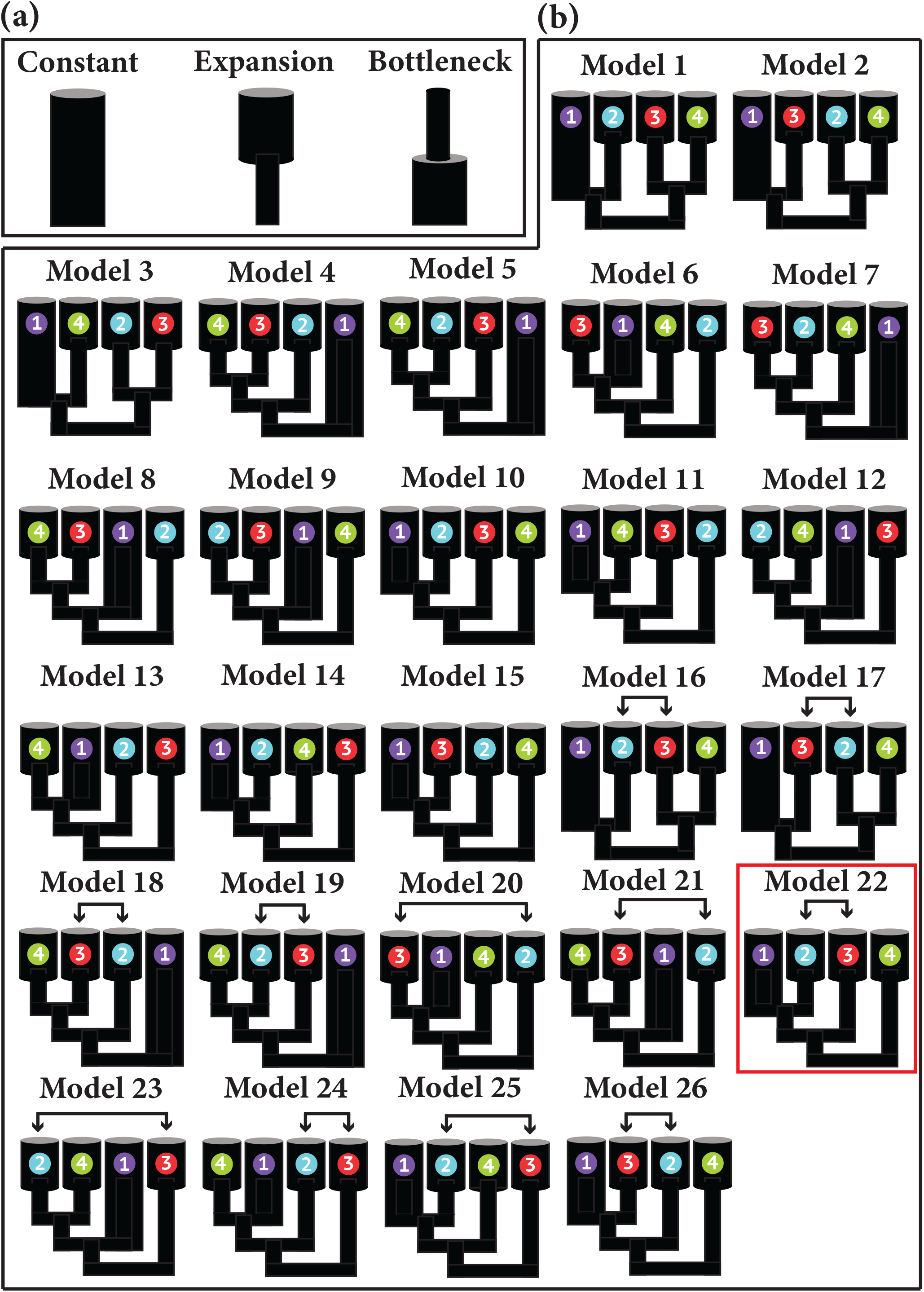
Representation of the models tested using convolutional neural networks. (a) set of three models used to test population trajectory through time; (b) set of 26 models used to test the evolutionary relationships and secondary contact of *Norops* ssp. Numbers and colors represent populations recovered in STRUCTURE analysis. Purple circle = *Norops brasilinesis* (population 1); blue circles = *N. brasilinesis* (population 2); red circles = *N. brasilinesis* (population 3); green circles = *Norops planiceps*. Gene between populations 2 and 3 is represented by arrows. The best-supported model for CNN in the second part of comparison is marked by a red box.

#### Testing evolutionary relationships and gene flow

In the second part, we implemented a CNN architecture to assess the relationships among populations and gene flow between populations that showed evidence of admixture in STRUCTURE. We specified 26 demographic models, which comprise the combination of all 15 possible topologies along with scenarios of isolation after divergence or secondary contact that reflect our identification of individuals that are potentially admixed. For example, because we recovered substantial admixture between populations 2 and 3, we included models with potential secondary contact between these populations (see Figure 4). We did not include models with secondary contact when populations 2 and 3 were sister in the phylogenetic tree, because it was impractical to distinguish between isolation and secondary contact models in our preliminary runs. We used fastsimcoal2 to generate 10,000 data examples per model. As in the first part, we generated short DNA sequences of 5 bp for 100,000 independent loci in a way to simulate 1 SNP per locus. However, we only output the number of SNPs observed in the empirical dataset. Parameters in these models include ancestral and current population size, the time of population size changing, divergence time, migration rate, time of migration, and topology. Priors are available in Table S1. We converted alignments nto images as described previously. In addition, because the relationship among populations is a key parameter in the models, images always presented populations in the same order: *N. brasiliensis* (population 1), *N. brasiliensis* (population 2), *N. brasiliensis* (population 3), and *N. planiceps*.

**Figure 4.**
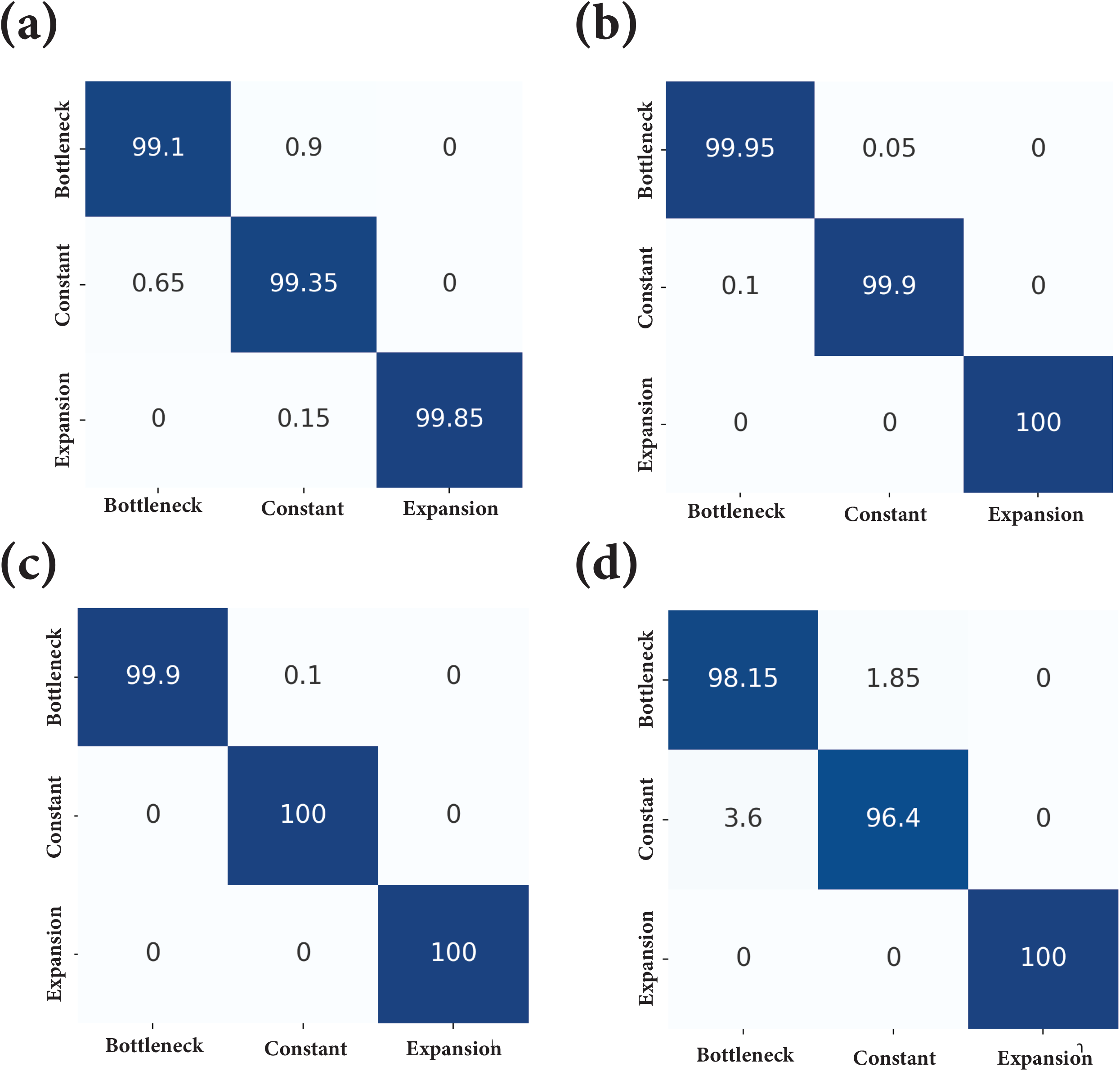
Confusion matrices measuring the accuracy of the trained CNNs model on the test dataset to detect demographic changes through time. Numbers represent percentages, which were calculated based on 2,000 images for each model. (a) *Norops brasilinesis* (population 1); (b) *N. brasilinesis* (population 2); *N. brasilinesis* (population 3); *N. planiceps*.

We used a simpler CNN architecture for the second part because it achieved a higher accuracy when compared to the CNN architecture used in the first part. We built the CNN using a two-dimensional convolution layer (kernel = 3 × 1), a two-dimensional maximum pooling layer (kernel = 3 × 1). After that, we flattened the output layer from the pooling and generated a fully connected layer with 500 neurons using the hyperbolic tangent function (tanh) for all layers, followed by our final layer with 26 neurons, corresponding to different models (Figure 5), where we used the softmax function. We compiled our model similar to the first part: Adam optimization and categorical cross-entropy loss function, but we used a mini-batch size of 50. We trained the CNN for 5 epochs; but the model did not improve after the second epoch. Then we split simulations in training (80%) and test datasets (20%). Finally, we used the trained model to predict the empirical dataset. We used the python library Keras throughout to build the CNN.

**Figure 5.**
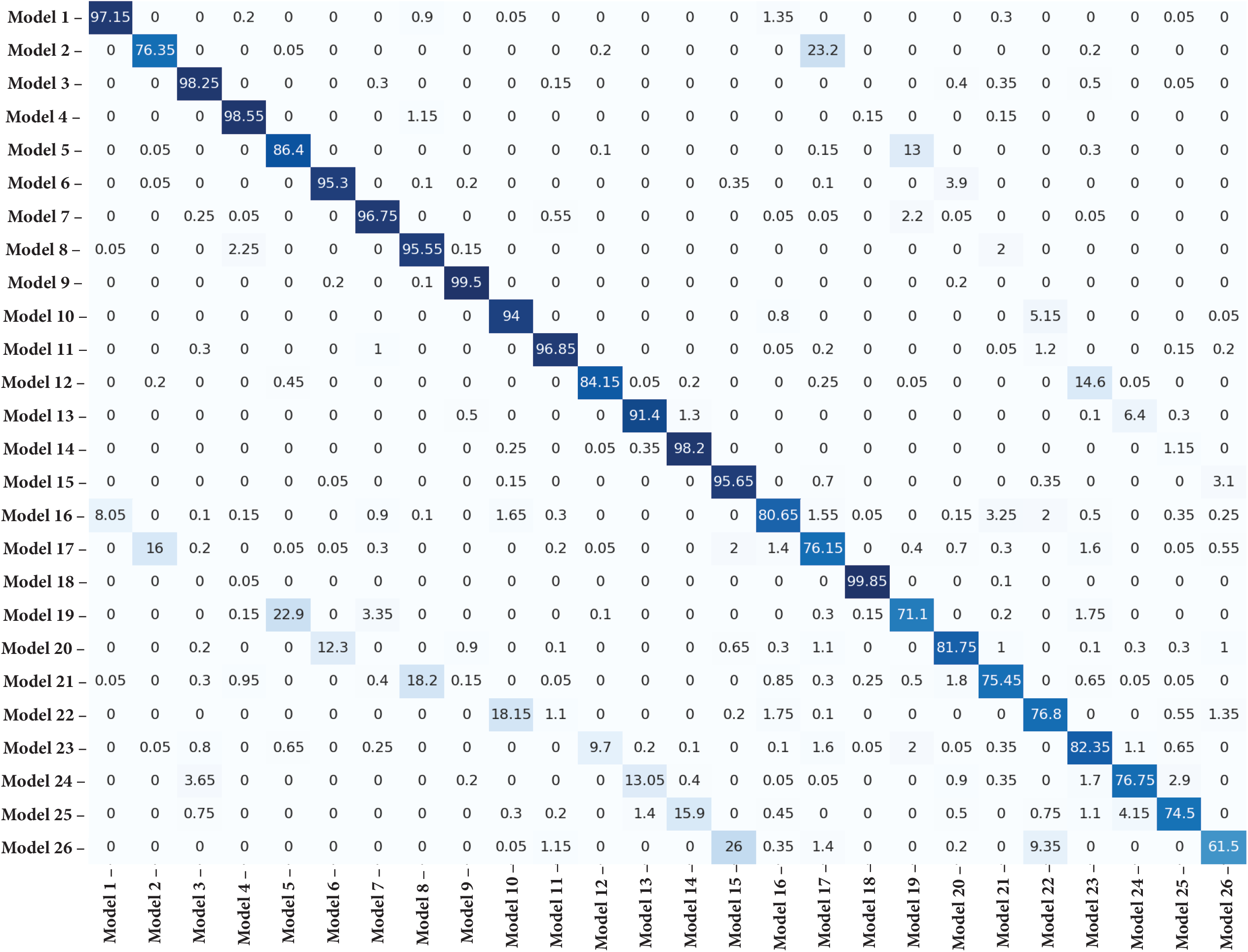
Confusion matrices measuring the accuracy of the trained CNNs model on the test dataset of 26 phylogeographic models. Numbers represent percentages, which were calculated based on 2,000 images for each model.

### Model selection in an approximate Bayesian computation framework

We also evaluated ABC performance for the second part of comparisons (from models 1 to 26). First, we used the R-package “PipeMaster” to perform 100,000 simulations for each model to generate summary statistics (Gehara et al. in prep.; www.github.com/gehara/PipeMaster). PipeMaster is a user-friendly R-package that builds demographic models and then simulates data under the coalescent process using *msABC* (Pavlidis et al., 2010). Demographic models mirrored empirical datasets with respect to the number of populations, the number of individuals within each population, and the number of loci. Priors used to build the models were the same used to construct CNNs models and are presented in Table S1. After simulations, we used the ABC approach to estimate model support using the “postpr” function implemented in “abc” R-package. We set the tolerance value to 1% and used the rejection method to compare models. We evaluate whether simulations produced summary statistics similar to the empirical dataset using PCAs.

## Results

### Genomic data processing

After genomic data processing, we obtained 4174 unlinked SNPs when all samples were combined, or 6860, 10931, 9396, and 12048 unlinked SNPs for the three *N. brasiliensis* populations and *N. planiceps*, respectively. Because our CNN approach does not accommodate missing data, loci were required to be present in 100% of the samples.

### Population assignment

The STRUCTURE analysis recovered four geographically structured populations that correspond to *N. planiceps* and three populations within *N. brasiliensis* (hereafter population 1, population 2, and population 3; Figure 2). While *N. planiceps* is distributed in northern Amazonia, population 1 is found in an enclave of Seasonally Dry Tropical Forests within Cerrado. Population 2 is more widespread in Cerrado and population 3 is found in lowlands within Cerrado. In addition, population assignment analysis revealed a region of high admixture between population 2 and 3 (locality #9).

### Demographic model selection

We recovered population expansion as the best demographic scenario for *N. planiceps*, population 2, and population 3 with a probability of 0.99, 0.59, and 1.0, respectively (Table 2). For population 2, the lower probability value is likely related to the unaccounted gene flow with populations 3, which introduced a genetic variation that was not captured by the model. Conversely, for population 1, we found evidence of constant population size over time (probability = 0.985; Table 2). For all models within each population, the CNN model had a high accuracy when predicting the test set labels, reaching an accuracy higher than 99% for all models (Figure 4).

**Table 2.** The probability of each model tested using convolutional neural networks and approximate Bayesian computation. Comparisons were first performed within part 1 only using CNNs, and subsequently, models in part 2 were constructed based on demographic scenario inferred in part 1. The best-fit model selected in each part is highlighted in bold.

| Part 1 |  | Part 2 |  |  |
| --- | --- | --- | --- | --- |
| Model | CNNs Probability | Model | CNNs probability | ABC posterior probability |
| **Population 1 - Constant** | **0.985** | Model 1 | 0 | 0.033 |
| Population 1 - Expansion | 0.015 | Model 2 | 0 | 0.035 |
| Population 1 - Bottleneck | 0 | Model 3 | 0 | 0.025 |
|  |  | Model 4 | 0 | 0.11 |
|  |  | Model 5 | 0 | 0.11 |
| Population 2 - Constant | 0.41 | Model 6 | 0 | 0 |
| **Population 2 - Expansion** | **0.59** | Model 7 | 0 | 0.1 |
| Population 2 - Bottleneck | 0 | Model 8 | 0 | 0.008 |
|  |  | Model 9 | 0 | 0.007 |
|  |  | Model 10 | 0.01 | 0 |
| Population 3 - Constant | 0 | Model 11 | 0 | 0 |
| **Population 3 - Expansion** | **1.0** | Model 12 | 0 | 0.007 |
| Population 3 - Bottleneck | 0 | Model 13 | 0 | 0 |
|  |  | Model 14 | 0 | 0 |
|  |  | Model 15 | 0 | 0 |
| *N. planiceps* - Constant | 0.01 | Model 16 | 0 | **0.15** |
| ***N. planiceps* - Expansion** | **0.99** | Model 17 | 0 | **0.15** |
| *N. planiceps* - Bottleneck | 0 | Model 18 | 0 | 0.037 |
|  |  | Model 19 | 0 | 0.038 |
|  |  | Model 20 | 0 | 0.0134 |
|  |  | Model 21 | 0 | 0.022 |
|  |  | Model 22 | **0.79** | 0.060 |
|  |  | Model 23 | 0 | 0.0046 |
|  |  | Model 24 | 0 | 0.012 |
|  |  | Model 25 | 0 | 0.013 |
|  |  | Model 26 | 0.20 | 0.065 |

For the second part of model comparison, CNN recovered a single model (#22) as the best evolutionary scenario with a probability of 0.79 (Table 2). As expected, *N. planiceps* was recovered as the sister species of *N. brasiliensis* and population 1 is more closely related to population 2 than to population 3. In addition, we found evidence of secondary contact between populations 2 and 3. The second-best model (model 26; probability = 0.20; Table 2) is similar to the best model but, in this scenario, population 1 is more closely related to population 3. All other scenarios had a probability of less than 1% (Table 2). Even comparing complex evolutionary histories, our CNN showed a high average accuracy: 87%; range: 62%–99%; Figure 5). Conversely, the posterior probabilities of ABC models were considerably lower. Models 16 and 17 were the best models supported by this analysis, with a posterior probability of 15% each (Table 2). PCAs showed that most models produced summary statistics coincident with empirical datasets, indicating that the choice of priors was plausible (Figure S1).

## Discussion

Our simulation testing implies that a deep learning approach for phylogeographic model selection can be very accurate for certain types of demographic processes. For example, the best CNN model had an accuracy of over 99% when testing for changes in effective population size through time in population 1 (i.e., constant, expansion, and bottleneck). We also found similar results for populations 2 and 3 (accuracy > 99%). Model accuracy was slightly lower for *N. planiceps*, likely caused by the small number of samples for this species. Model accuracy, therefore, seems to rely on the number of individuals and the number of SNPs. Even though we generated fewer SNPs for population 1, this model achieved higher accuracy than the one for *N. placenips* probably because we had twice the number of samples for population 1. For models 1 to 26, the average accuracy was 87%. Undoubtedly, these models are more complex than those dealing only with changes in population size, given that all populations were compared simultaneously, and we also included the relationships among them, gene flow between populations 2 and 3, and divergence times. Still, our approach reached an accuracy similar to other approaches. Conversely, ABC was unable to accurately find the best fit model, given the low posterior probability of all models (see Table 2).

CNN and ABC share many similarities, including the use of a simulation-based approach to generate new examples, given a demographic scenario and a set of priors. However, one of the main differences between these approaches is how they summarize the simulated datasets and, most importantly, how empirical and simulated datasets are compared. Therefore, a key feature of any of these methods is to be able to summarize the information in the data in a meaningful way. For ABC, a large number of summary statistics is usually calculated from the simulated datasets, e.g., Tajima’s D, nucleotide diversity, F_ST_, and Fu and Li’s D and F statistics. Each summary statistic has been used in phylogeographic investigations. For example, Tajima’s D is a summary statistic that detects departures from constant population sizes over time, including population expansion and bottleneck. In addition, fixation indexes have measured the degree of differentiation among populations. The choice of summary statistics is largely subjective, with most studies choosing not to identify a subset of summary statistics that maximize model probability. As stated by Beaumont et al. (2002), “*a crucial limitation of the rejection-sampling method is that only a small number of summary statistics can usually be handled*”. Our results mirror those from previous research suggesting that ABC does not perform as well with large numbers of models (Pelletier & Carstens, 2014; Smith et al., 2017).

Although it is beyond the scope of this study to compare different methods of phylogeographic model selection, a broadly comparison of the accuracy of these approaches can be made based on our approach. For example, PHRAPL summarizes data using gene trees and because of that, incomplete lineage sorting (ILS) is one of the main sources of model selection inaccuracy (Jackson et al., 2017). At shallower divergence times, a more pronounced discordance in gene trees can be observed and, consequently, it is more difficult to identify the evolutionary scenario that gives birth to the data. Similarly, for CNNs, as the divergence times decreases among lineages the model accuracy decreases, which likely results from ILS (Blischak et al., 2020). As noted above, conventional ABC approaches can attain high accuracy with a high number of models, but this potential liability can be alleviated. For example, Smith et al. (2017) proposed a Random Forest approach to test 15 evolutionary scenarios for a land snail endemic to the Pacific Northwest of North America and compare the Random Forest classifier with ABC. Their overall errors using Random Forest were 7.67% (range: 0–42%) and ∼30% for ABC. The overall error of our CNN was 13%, but we noticed that most misclassification was between models that only differed on the presence or absence of secondary contact. Since Smith et al. (2017) did not include gene flow in the tested models, we subset our models and trained a CNN only with isolation models (models 1 to 15). The overall error was 1.5% (0.75–3%; Figure S2) and the best model had a probability of 87%. In a more recent study, Smith & Carstens (2020) applied Random Forest to the reticulate taildropper slug (*Prophysaon andersoni*) and found an average error of 5.2% when comparing 208 demographic models. These results show that CNN has an accuracy comparable to the best results reported for other methods (i.e., ABC with Random Forest). Unfortunately, the comparison between CNN and AIC-based methods (such as PHRAPL) is not straightforward because they use different frameworks to measure model performance. In particular, AIC-based approaches to model selection lack the built-in approach for assessing model accuracy (i.e., identifiability) that deep learning approaches such as CNN and ABC with Random Forest include.

One advantage of CNNs is that researchers are absolved of the requirement to summarize their data using summary statistics. Since there exists a set of statistics that is likely best used with a particular demographic history, this is particularly challenging for investigations into non-model systems. In our system (*N. planiceps* and *N. brasiliensis*) and others, there is a scarcity of *a priori* ecological and evolutionary information that limits the ability of researchers to specify a small set of candidate models and choose appropriate summary statistics. In such a scenario, approaches such as CNNs, PHRAPL, and delimitR offer the potential to compare among a large number of competing alternatives models without the need to make choices that are likely to influence the outcome. That is not to say that CNN approaches require the data to be summarized as we have done here. For example, Blischak et al. (2020) used CNNs to detect hybridization in simulated and an empirical system from *Heliconius* butterflies. They simulated chromosome-scale data for four species and generated images based on the pairwise Nei’s genetic distance among populations. Their approach was found to be more accurate than approaches that were based on introgression-specific summary statistics.

Our approach was computationally more demanding than the one proposed by Blischak et al. (2020). It requires an average of two seconds to run the simulation in *fastsimcoal2* and eight seconds to process the image (∼ 10 seconds from simulation to generate an image). Since we simulated 10,000 examples per model, it would take about 27 hours to simulate the images that correspond to one scenario. It required 10 hours to run one epoch in the comparison among 26 models (208,000 training images and 52,000 test images), but this time can be optimized by using Graphical Processing Unit (GPU) instead of Central Processing Unit (CPU). Although the simulation and CNN were performed using the resources provided by the Ohio Supercomputer Center, we used a Mac mini (1.6 GHz Intel Core i5, 8 GB RAM, 2 cores) to generate these reference values to provide context for potential users of this approach who do not have access to supercomputing centers. By far the biggest computational hurdle was the number of images storage in the Supercomputer. Our analysis used a total of 380,000 images.

### Evolutionary history of South American lizards

Pleistocene climate change has been proposed as one of the main drivers of speciation at higher latitudes (Burbrink et al., 2016; Hewitt, 2000, 2004). The Pleistocene refugia hypothesis (PRH) posits that species had to inhabit favorable refugia to persist and thrive under the new environmental conditions (Vanzolini & Williams, 1970). In South America, Haffer (1969) and Vanzolini & Williams (1970) almost simultaneously proposed the PRH to explain patterns of species diversity and distribution in the Amazon rainforest, where climate oscillations led to a series of contraction events of rainforests and expansions of dry vegetations during glacial periods, which would enable allopatric speciation of the associated biota. While this has been a popular hypothesis, many investigations have dismissed the Pleistocene refugia model based on multiple biological and paleoenvironmental sources of evidence (Bush & Oliveira, 2006; Lessa et al., 1997; Thomé et al., 2010; Wang et al., 2017). Cheng et al. (2013), based on speleothem oxygen isotope records, proposed an alternative speciation model for the Late Pleistocene in South America, in which a quasi-dipolar precipitation pattern during the Pleistocene would respond for differences in biodiversity between western and eastern Amazonia. In particular, eastern Amazon, which is more connected to the historical and current climate in the Cerrado, held desynchronized interleaved periods of wet and dry climates during the last 250 thousand years (kyr) with western Amazonia. These climatic patterns resulted in habitat fragmentation that isolated species that were previously broadly distributed and led to decreased gene flow and increased genetic differentiation. Some community-level analyses suggest that this model is broadly applicable (Gehara et al., 2017; Silva et al., 2019). In contrast, climate was more stable in western Amazon, which is hypothesized to have generated the observed higher levels of biodiversity across multiple taxonomic groups and likely population stability through time.

Our phylogeographic model selection results support the quasi-dipolar scenario of Cheng et al. (2013). We found support for population expansion in *N. planiceps* and populations 2 and 3. While our results showed that population 1 was constant in size through time, this population is located in an enclave of Caatinga within Cerrado (Paranã valley). Caatinga is the largest nucleus of Seasonally Dry Tropical Forests (SDTF) and characterized by xeric vegetation, high seasonality, and unpredictable droughts. It is hypothesized that the climatic oscillations during the Pleistocene led the expansion and connection of now disjunct SDTFs (the Pleistocenic Arc Hypothesis - PAH; Prado & Gibbs, 1993; Pennington et al., 2000). This hypothesis is supported by the disjunct distribution of plants and animals as well as molecular data (Lanna et al., 2018; Pennington et al., 2000; Werneck & Colli, 2006). However, the exact time of the PAH is uncertain and the SDTFs could have expanded earlier, during the transition between Pliocene and Pleistocene, and have fragmented before the Last Glacial Maximum (Werneck et al., 2011), which could explain the stable population sizes we recovered in the longer term.

In addition to climatic oscillations, the pattern of diversification found by our study mirrors the current taxonomic status of both species, though we found a hidden genetic diversity within *N. brasiliensis*. The pattern of divergence among lineages within *N. brasiliensis* follows a southeast-northwest pattern of differentiation, which is shared with other squamates in Cerrado (Guarnizo et al., 2016; Prado et al., 2012; Santos et al., 2014). This pattern of differentiation was likely driven by landscape features and climatic conditions.

### Conclusion

Deep learning techniques have been successfully used in fields like medical sciences and agriculture, but their usage in evolutionary biology has just begun (but see Blischak et al., 2020; Flagel et al., 2019; Schrider & Kern, 2018). Our results showed that CNNs can be an effective and promising approach for phylogeographic model selection. We showed that a DNA alignment can be used as the source of comparison of a large number of models, without the need of genetic summary statistics. Also, our approach revealed a complex evolutionary scenario among lizards distributed in contrasting environments in South America, which involves hidden genetic diversity, gene flow between non-sister populations, and changes in effective population size through time. Finally, we encourage future investigations to compare the relative performance of different approaches for phylogeographic model selection.

## Acknowledgements

We thank members of the Carstens Lab for comments and suggestions on the manuscript and Lisa N. Barrow and Megan L. Smith for lab assistance. We thank Ohio Supercomputer Center for computing resources. We thank curators and managers of the INPA-H and INPA-HT (C. Ribas, M. Freitas, A. Silva) for granting and processing samples under their care. EMF thanks the Coordenação de Aperfeiçoamento de Pessoal de Nível Superior (CAPES) for his doctoral fellowship (process #88881.170016/2018). FPW thanks Conselho Nacional de Desenvolvimento Científico e Tecnológico (CNPq) for her productivity fellowship (process #305535/2017-0). GRC thanks CAPES, CNPq, Fundação de Apoio à Pesquisa do Distrito Federal – FAPDF and the USAID’s PEER program under cooperative agreement AID-OAA-A-11-00012 for financial support.

## Author Contributions

EMF and BCC conceived the ideas and designed methodology; EMF conducted the lab work and conducted the analyses. All authors interpreted the results and participated in the writing of the manuscript and gave final approval for submission.

