## Supplemental table and figures for "Phylogeographic model selection using convolutional neural networks"

### #

### *Molecular Ecology Resources*

Supporting Information

\#

**Table S1.** Information on priors used in this study. Population sizes represent the numbers of haploid individuals. Divergences times and changes in the effective population sizes are in generations before the present. Changes in the effective population sizes and migration rates were parameterized according fastsimcoal2 manual. All priors were sampled from a uniform distribution. Populations from 1 to 3 refer to the recovered populations in STRUCTURE analysis (see results).

| **Priors** | **Lower bound** | **Upper bound** |
| --- | --- | --- |
| Current population size (Population 1) | 50,000 | 200,000 |
| Current population size (Population 2) | 50,000 | 200,000 |
| Current population size (Population 3) | 50,000 | 200,000 |
| Current population size (*Norops planiceps*) | 50,000 | 200,000 |
| Ancestral population size (Population 1) | 1,000 | 10,000 |
| Ancestral population size (Population 2) | 1,000 | 10,000 |
| Ancestral population size (Population 3) | 1,000 | 10,000 |
| Ancestral population size (*Norops planiceps*) | 1,000 | 10,000 |
| Expansion or bottleneck time (Population 1) | 5,000 | 21,000 |
| Expansion or bottleneck time (Population 2) | 5,000 | 21,000 |
| Expansion or bottleneck time (Population 3) | 5,000 | 21,000 |
| Expansion or bottleneck time (*Norops planiceps*) | 5,000 | 21,000 |
| Divergence time (T1) | 5,000 | 20,000 |
| Divergence time (T2) | 25,000 | 35,000 |
| Divergence time (T3) | 40,000 | 50,000 |
| Migration rates | 5 | 10 |

**Figure S1.** Principal Components Analysis plots (PC1 X PC2) derived from the simulated distributions of summary statistics (Models 1 to 26) and observed dataset.

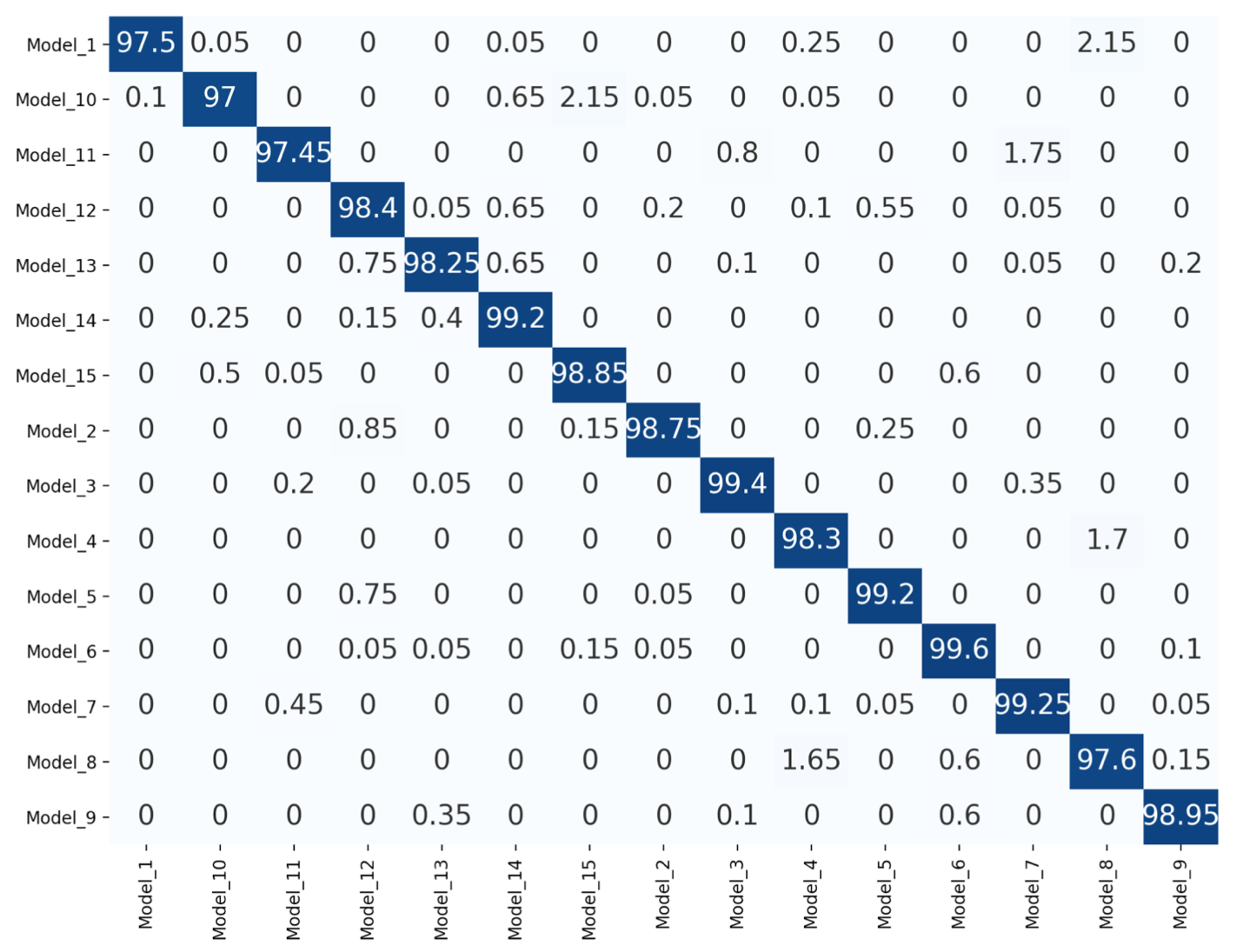

**Figure S2.** Confusion matrices measuring the accuracy of the trained CNNs model on the test dataset of only isolation models (from models 1 to 15). Numbers represent percentages, which were calculated based on 2,000 images for each model.
